# Modeling a co-culture of *Clostridium autoethanogenum* and *Clostridium kluyveri* to increase syngas conversion to medium-chain fatty-acids

**DOI:** 10.1101/2020.06.23.167189

**Authors:** Sara Benito-Vaquerizo, Martijn Diender, Ivette Parera Olm, Vitor Martins dos Santos, Peter J. Schaap, Diana Z. Sousa, Maria Suarez-Diez

## Abstract

Microbial fermentation of synthesis gas (syngas) is becoming more attractive for sustainable production of commodity chemicals. To date, syngas fermentation focuses mainly on the use of *Clostridium* species for the production of small organic molecules such as ethanol and acetate. The cocultivation of syngas-fermenting microorganisms with chain-elongating bacteria can expand the range of possible products, allowing, for instance, the production of medium-chain fatty acids (MCFA) and alcohols from syngas. To explore these possibilities, we report herein a genome-scale, constraint-based metabolic model to describe growth of a co-culture of *Clostridium autoethanogenum* and *Clostridium kluyveri* on syngas for the production of valuable compounds. Community flux balance analysis was used to gain insight into the metabolism of the two strains and their interactions, and to reveal potential strategies enabling production of butyrate and hexanoate. The model suggests that addition of succinate is one strategy to optimize the production of medium-chain fatty-acids from syngas with this co-culture. According to the predictions, addition of succinate increases the pool of crotonyl-CoA and the ethanol/acetate uptake ratio in *C. kluyveri*, resulting in the flux of up to 60% of electrons into hexanoate. Other potential way to optimize butyrate and hexanoate is to increase ethanol production by *C. autoethanogenum*. Deletion of either formate transport, acetaldehyde dehydrogenase or formate dehydrogenase (ferredoxin) from the metabolic model of *C. autoethanogenum* leads to a (potential) increase in ethanol production up to 150%, which is clearly very attractive.

## 1. Introduction

One of the biggest challenges society faces nowadays is finding alternative processes for the sustainable production of fuels and chemicals. At present, the production of many commodities depends on fossil fuels (not sustainable) or sugar crops (competing with human and animal food consumption) [1]. To circumvent this, circular approaches are required, such as the conversion of lignocellulosic biomass or municipal waste as feedstocks to fuels and chemicals [2]. Although lignocellulosic biomass has been identified as a promising source for renewable energy and carbon [3], current technologies involving hydrolysis of this substrate result in a complex mixture of compounds that need further separation and individual processing [4]. However, gasification of these rigid materials allows for the conversion of the carbon in the original source to synthesis gas (syngas), consisting mainly of CO, H_2_ and CO_2_. This energy-rich syngas can be further used as feedstock for chemocatalytic processes such as Fischer-Tropsh, but microbial fermentation of syngas is gaining more attention recently as a potential production platform [5], [6]. Compared to chemical catalysts, microorganisms are more robust to variations of CO/H_2_ ratio in syngas, and are also more resistant to the presence of certain impurities (e.g. sulfides), reducing the need for costly pre-treatment of syngas [7].

Acetogenic clostridia are efficient microbial hosts for syngas fermentation as they can grow on CO and CO/H_2_ via the Wood–Ljungdahl pathway [8]. However, the natural product range of most acetogens is limited to a mixture of acetate and ethanol [9]. Co-cultivation of a syngas-fermenting organism with other organisms (that use the primary products of syngas fermentation) can be used to extend the range of possible products. Previously, a co-culture of *Clostridium autoethanogenum* and *Clostridium kluyveri* was described to produce medium-chain fatty acids (C4-C6) and their respective alcohols by assimilation of CO or syngas [10, 11]. *C. autoethanogenum* is an acetogenic bacterium able to produce acetate and ethanol when growing on CO or syngas [12]. *C. kluyveri* grows on acetate and ethanol via reverse-*β*-oxidation, producing chain-elongated acids like butyrate and hexanoate.

When *C. kluyveri* is grown in co-culture with *C. autoethanogenum* on CO, it produces butyrate and hexanoate, which are further reduced by the acetogen to the corresponding alcohols, butanol and hexanol [10]. MCFA are used to produce pharmaceutical and personal care products, animal feed additives and lubricants, among other, and can be converted chemically or enzymatically into valuable biofuel molecules such as methyl esters, methyl ketones, alkenes and alkanes [13, 14]. The theoretical maximum yield of hexanoate production in a co-culture of *C. autoethanogenum* and *C. kluyveri* is 0.056 mmol of hexanoate per mmol of CO, whereas the yield obtained in the most recent study [11] was 0.009 mmol hexanoate per mmol of CO, so there is substantial room for improvement and new strategies need to be developed.

Genome-scale, constraint-based metabolic models (GEMs) attempt to represent the complete set of reactions in a living organism, and have been used to gain better understanding of cellular metabolism, assessing theoretical capabilities or designing media and processes [15]. GEMs can be used to link the microbial consumption and production rates with cellular growth rates. Moreover, they enable linking these phenotypes with the genome content of the studied organisms and with internal phenotypes, such as metabolic fluxes that are usually difficult to measure experimentally. GEMs and their analysis with constraint-based techniques, such as flux balance analysis (FBA) for the calculation of steady-states, have been proven effective tools to devise strategies for increasing productivity of microbial fermentation processes [16, 17, 18]. Specifically, GEMs have been used to further understand the metabolism of clostridia. For instance, the GEM of *Clostridium thermocellum* allowed the design of metabolic strategies to increase ethanol production after identification of bottlenecks in central carbon metabolism that were inhibiting its production [19]. Stolyar and collaborators ([20]), generated a multi-species GEM by combining the GEMs of bacterium *Desulfovibrio* and archeon *Methanococcus maripaludis* S2 into a single model with a shared extracellular environment, bringing the use of GEMs to a next level. Since then, this type of community models have been used to describe metabolic interactions among community members and inter-species fluxes [21]. Li and Henson [22], recently used GEMs to compare mono-culture and co-culture systems to produce butyrate from carbon monoxide. They applied dynamic flux balance analysis (dFBA) [23] to analyze a community GEM to cover the changes in community composition over time and to assess the relative performance of these mixed cultures. The availability of GEMs for *C. autoethanogenum* [24] and *C. kluyveri* [25], enables the use of community modeling as a potential method to help optimizing the performance of this co-culture for syngas fermentation to elongated acids and alcohols.

In this study, we present a multi-species model built by combining the GEMs of *C. autoethanogenum* [24] and *C. kluyveri* [25]. The model accounts for experimental measurements informing on relative species abundances and volumetric production rates of syngas fermentation products obtained in chemostat runs under different conditions [11]. In order to test the model, experimental values were introduced as environmental constraints by employing community flux balance analysis (cFBA) [21, 26]. Subsequently, the model was used to identify and assess strategies to optimize desired products, specifically butyrate and hexanoate.

## 2. Materials and methods

### 2.1. GEMs of C.autoethanogenum and C. kluyveri

To represent the metabolism of *C. autoethanogenum*, the previously described GEM, iCLAU786, was retrieved in SBML (XML) format from the supplementary material provided by Valgepea et al. [24]. This model was amended with an exchange reaction to simulate acetate uptake when this is used as additional substrate ((EX AC c). eQUILIBRATOR [27] was used to manually verify reaction directionality: Gibbs energy released (Δ*G*) at pH 7.0 and ionic strength (0.1M) was computed. Reactions with Δ*G ∈* [−30, 30] kJ/mol were considered reversible.

The GEM of *C.kluyveri*, iCKL708, was downloaded in table format from its publication [25]. An additional reaction was added to excrete biomass, which was first included as new metabolite and as additional product in the biomass reaction BOF. Minor changes were applied affecting the reversibility of few reactions and addition of protons. Pyruvate synthase (Rckl119) was set to non-reversible in the direction of pyruvate production [28]. Pyruvate formate lyase (PFL) was set to non-reversible, allowing only the production of formate and acetyl-CoA. Protons were added in the exchange of heptanoate reaction (Rckl870). eQUILIBRATOR [27] was used to manually verify reaction directionality [27] as in previous model. The updated model was converted to SBML level 3 version 1 standardization [29].

### 2.2. Multi-species GEM reconstruction

The multi-species GEM of *C. autoethanogenum* and *C. kluyveri* was generated by combination of single species models: iCLAU786 [24] and the updated version of iCKL708 [25], respectively, following a compartmentalized approach [20] were each species is considered a single compartment. Therefore, we consider two internal compartments: ‘cytosol_auto’ and ‘cytosol_kluy’, with ‘c’ and ‘ck’ as their respective identifier (id). Intracellular metabolites were assigned to their corresponding compartment and the flag ‘_c’ was added to the id of metabolites belonging to ‘cytosol auto’ and ‘_ck’ to those belonging to ‘cytosol_kluy’. Metabolites that can be exchanged between species, are also defined in a common extracellular compartment, by using the “_e” tag. This means, that all extracellular metabolites need to have the same naming system (name-space) and be unique for both species.

Each species has its own biomass synthesis reaction. An extra biomass metabolite was created and defined in the extracellular compartment for each species: ‘BIOMASS_c_*e*’ and ‘BIOMASS_ck_*e*’. In addition, two extra reactions were added for each species, one to transport biomass from the intracellular to the extracellular compartment, and a second one to excrete biomass (exchange reaction). A reaction was included to distinguish the amount of H_2_ excreted by *C. kluyveri*, from the amount of H_2_ metabolized by *C. autoethanogenum*. The same was done for acetate. A reaction was included to distinguish the amount of acetate metabolized by C. autoethanogenum, from the amount of acetate produced by *C. autoethanogenum*. The model also contemplates the possible production of butanol and hexanol via butyrate and hexanoate uptake by *C. autoethanogenum*. The added reactions are a transport reaction from the external compartment to the internal compartment of *C. autoethanogenum*, reactions for production of butyraldehyde and caproaldehyde from the corresponding fatty-acids and reactions for production of alcohols from their corresponding aldehydes. Finally, the multi-species model was transformed into SBML level 3 version 1 (see supplementary material).

### 2.3. Multi-species modeling framework

In order to model the community, we have followed an approach similar to the one proposed by SteadyCom [30] and that is based on community FBA (cFBA) [26]. Environmental fluxes (mmol l^−1^ h^−1^) are integrated as model constraints instead of specific fluxes (mmol gDW^−1^ h^−1^), where gDW indicates grams of dry weight. The biomass reaction of each species incorporates as new term, the biomass of the relative species together with the growth rate term. In this way, we can account for species abundance in the community. The biomass of each species is calculated based on the community biomass and the species ratio. In addition to this, steady-state and equal growth rate of species are assumed.

### 2.4. Calculation of species abundances

The ratio between *C. autoethanogenum* and *C. kluyveri* in co-culture, was estimated from transcriptomics reads and from dry weight calculations based on microscopy observations.

Transcriptomic data was obtained from steady-state co-cultures grown in chemostats [11]. The Genomes of *C. autoethanogenum*: DSM 10061 (GCA 000484505.1) [31] and *C. kluyveri* : DSM 555 (GCA 000016505.1) [32] were retrieved from the European Nucleotide Archive. The genomes have similar size with sequence length 4.352.205 and 4.023.800, respectively [33]. Reads were mapped to each genome using BWA-SW (Burrows Wheeler Aligner) [34] and the ratio was calculated based on the amount of reads associated to each species.

The cellular volume of each species was calculated based on their average size. *C. autoethanogenum* is a rod-shaped bacterium with an average size of 0.5 × 3.2 µm [12] and *C. kluyveri* cells are curved rods, with average size of 12.5 µm in length and 1.5 µm in width [35]. Cell volume was calculated following a previously proposed formula for rod-shaped cells [36]: *V* = [(*w*^2^· *π/*4) · (*l* − *w*)] + (*π* · *w*^3^*/*6)], with *l* and *w* indicating length and width respectively. The associated dry weight (DW) was then derived using: *DW* = 435 · *V* ^0.86^ [36]. A proportion observed in microscopic images between cell numbers (10 *C. autoethanogenum* by 1 *C. kluyveri*) was taking into account to calculate the accumulated dry weight. So, the dry weight of *C. autoethanogenum* was multiplied by 10 and the dry weight of *C. kluyveri* was multiplied by 1. Finally, the biomass-species ratio was calculated based on the ratio of the accumulated dry weights. The proportion was observed to be constant among all experimental conditions.

### 2.5. Use of experimental values to constrain the model

Experimental measurements were converted to mmol l^−1^ h^−1^. Product concentrations, measured in mM, were used to compute product secretion rates (mmol l^−1^ h^−1^) by using the set hydraulic retention time (HRT) of the chemostats. Growth rate was calculated as the inverse of the HRT and expressed in h^−1^. In co-culture, growth rate of both species was assumed to be the same and equal to that of the community, since HRT was kept constant both, in mono-culture and co-culture experiments. We have followed the usual convention in constraint-based modeling, so that uptake is represented by negative fluxes whereas production corresponds to positive fluxes. To model experimental conditions, we fix substrate uptake rates to the desired ones by setting the lower bounds of the corresponding exchange reactions to the measured values multiplied by −1 (as it corresponds to consumption). Biomass reactions were constrained with the growth rate multiplying the total biomass by the ratio of each species (gDW l^−1^ h^−1^). Similarly, product formation was set to be at least 80% of the calculated product formation by modifying the lower bound of the corresponding exchange reaction. ATP maintenance reactions of each species, ATPM auto and ATPM, were transformed to mmol l^-1^ h^-1^ from the pre-set values in mmol gDW^−1^ h^−1^ multiplying by the total biomass and species ratio. In cases where metabolites behave as products that are further assimilated by the other species, the transport reactions of these metabolites are forced to operate in the direction from the external compartment to the other species compartment.

#### 2.5.1. Chemostat experimental data

Experimental data was collected from reactor run 3 and 4 of the recent study [11] on *C. autoethanogenum* in mono-culture and co-cultivation of *C. autoethanogenum* and *C. kluyveri* grown on CO/H_2_ and CO/acetate. In coculture experiments, *C. kluyveri* was inoculated in the reactor on top of *C. autoethanogenum* in a 1:20 volume ratio. The organisms were cultivated in chemostat to control environmental conditions such as pH (6.2), temperature (37 °), HRT (between 1.5 to 2 days) and medium composition. Total reactor volume is 1.5 l. Working volume was set between 0.75 l to 1 l. Experiments were run on different conditions of CO/H_2_ and CO/acetate as initial substrates. Concentrations of organic acids and alcohols in the reactor and gas composition in the outflow were tracked during the runs.

### 2.6. Model simulations

Model simulations were done using COBRApy, version 0.17.0 [37], IBM ILOG CPLEX 128 and Python 3.6. Simulations based on changes/addition of parameters were done by constraining the associated reactions with the mentioned values. When simulations required removal of a substrate or product, flux through the associated reaction was set to 0. Constraints on the profile of fermentation products were kept unchanged when simulations were based on substrate uptake ratios in *C. kluyveri*, unless stated otherwise. For each explored condition, the solution space and the set of fluxes compatible with the measured constraints was sampled using the *sample* function in the flux analysis submodule COBRApy. Flux sampling is a method to get a distribution of fluxes [38] under specific conditions. Presented results are the average and standard deviation based on 15000 iterations generated at each condition. All additional assumptions taken into account during model simulations are listed in the supplementary material.

### 2.7. Genetic intervention strategy

OptKnock and RobustKnock [39, 40] were applied as algorithms that suggest reactions to be knocked out that can potentially increase the yield of a target reaction. The algorithms were applied to increase ethanol production in the GEM of *C. autoethanogenum*. Both algorithms were integrated in a python script adapted for COBRApy and CPLEX as solver. OptKnock identifies a set of reaction knockouts that allows high production of a target product under the constraint of optimal growth in wild type. RobustKnock guarantees a minimal production rate by considering alternative optimal solutions that produce less of the target product. This is achieved by employing a bi-level max-min optimization. The possible reactions to be modified were adjusted in order to avoid essential reactions, reactions associated to essential genes, extracellular reactions and reactions with no associated genes.

## 3. Results

The objective of this study is to find optimization strategies for the production of medium-chain fatty-acids from syngas using the co-culture of *C. autoethanogenum* and *C. kluyveri*. The generated multi-species GEM, together with the GEM of *C. autoethanogenum*, were used to assess these strategies.

### 3.1. Description and validation of the GEM of individual strains

The GEM of *C. autoethanogenum*, iCLAU786 is composed of 1108 reactions and 1094 metabolites. The model is able to simulate growth on CO or syngas as the sole carbon and energy source, producing acetate and ethanol as the main fermentation products. According to model predictions, *C. autoethanogenum* is not able to produce medium-chain fatty-acids, which is in accordance with experimental observations [10].

The updated GEM of *C. kluyveri*, iCKL708 has 993 reactions and 811 metabolites. The model simulates growth on acetate and ethanol along with CO_2_ assimilation producing butyrate and hexanoate as the main chain-elongated products and H_2_.

### 3.2. Multi-species GEM

The multi-species GEM contains 2064 reactions and 1823 metabolites, from which 139 reactions correspond to extracellular reactions and 208 metabolites belong to the shared extracellular compartment. Figure 1 shows the dependencies included in the model to describe the syngas fermentation process by the co-culture. H_2_ ck reaction represents the amount of H_2_ excreted by *C. kluyveri*. H_2__e ← represents the uptake of H_2_ in *C. autoethanogenum*. The reaction ← AC_c, represents the uptake of acetate by *C. autoethanogenum* in simulations where acetate acts as additional substrate. This serves to distinguish the fluxes between acetate feed rate, acetate production rate and acetate shared between species. These are special cases since H_2_ and acetate can be shared, metabolized and produced in co-culture conditions.

**Figure 1:**
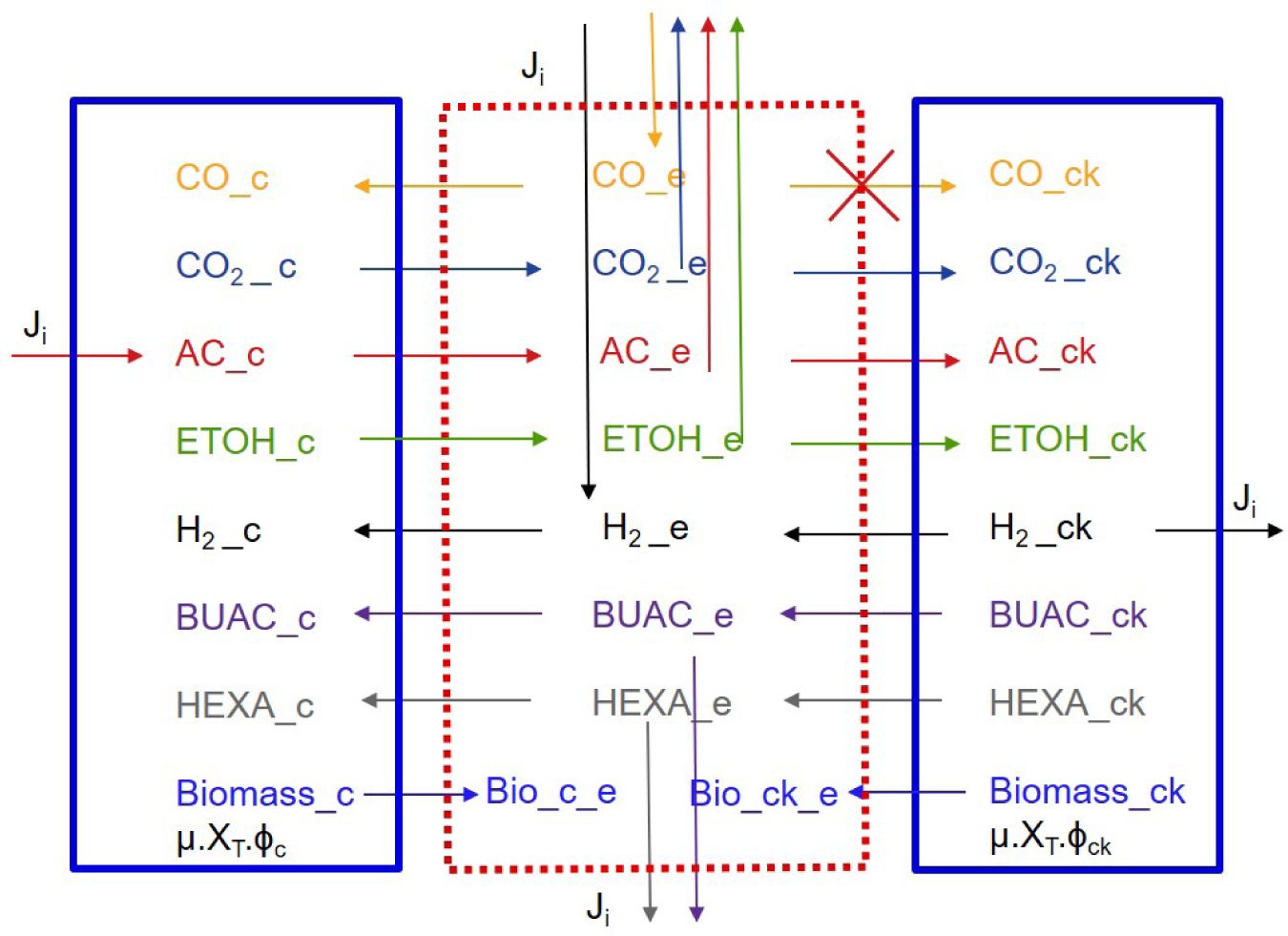
Dependencies applied to the multi-species model to describe possible interactions. Metabolites on the left belong to *C. autoethanogenum*’s compartment and on the right, to *C. kluyveri* ‘s compartment. Metabolites in the middle correspond to the extra-cellular compartment. Arrows between metabolites indicate transport reactions of that metabolite from one species’ compartment to the extracellular compartment or the other way around. Arrows affecting single metabolites indicate uptake or production of that metabolite. CO: carbon monoxide; H_2_: Hydrogen; AC: acetate; ETOH: ethanol; BUAC: butyrate; HEXA: hexanoate; J_i_: environmental fluxes of reaction i; µ: growth rate; X_T_: community biomass; ϕ_i_: species abundance, with *i* equals c or ck for *C. autoethanogenum* or *C. kluyveri*, respectively.

Previous studies have shown that *C. autoethanogenum* is able to grow on CO, CO/H_2_ producing ethanol and acetate as the main fermentation products [12]. Acetate and ethanol can further be taken up by *C. kluyveri* producing H_2_, butyrate and hexanoate. H_2_ produced by *C. kluyveri* appears to be further metabolized by *C. autoethanogenum* [10, 11]. Furthermore, the presence of aldehyde ferredoxin oxidoreductase and Ethanol:NAD+ oxidoreductase enzymes in *C. autoethanogemum* allows for a potential two step conversion of butyrate and hexanoate, via the respective aldehdye, to butanol and hexanol respectively. Previous experimental trials [10] showed that CO is not metabolized by *C. kluyveri* so, in the model, we prevented flux through the reaction *CO*_*e*_ → *CO*_*ck*_, as indicated in figure 1.

Microscopy observation of the co-culture led to the estimation of a ratio of 10 cells of *C. autoethanogenum* per 1 cell of *C. kluyveri*. Analyses of transcriptome samples obtained by RNAseq of the community [11], were done to identify the fraction of RNA arising from each community member. The estimated relative abundances yielded between 90-95% of *C. autoethanogenum* and 5-10% of *C. kluyveri*. Differences in cell size and volume were considered as *C. kluyveri* cells have approximately 36 more volume that *C. autoethanogenum* [12, 35, 36]. The estimated volumes were used to estimate dry weight of each cell species, resulting the cell dry mass of *C. kluyveri*, 22 times more than *C. autoethanogenum*. Finally, the cell ratio (10:1) was taken into account resulting in a biomass ratio of 68.5% *C. kluyveri* and 31.5% *C. autoethanogenum*. This cell ratio was observed in all experimental conditions and therefore, it was maintained constant in all model simulations.

### 3.3. Multi-species GEM accurately predicts experimental results

The initial mono-culture experiments only involved *C. autoethanogenum* [11]. Experiments were run on CO/H_2_ and CO/acetate as initial substrates. Figure 2 shows that the model predictions match relatively well the experimental results for *C. autoethanogenum*. Accordingly, the model predicts correctly that ethanol production increases gradually with the addition of H_2_ or acetate. Also, the model predicts no production of medium-chain fatty-acid in mono-culture, which agrees with experimental observations. However, the model predicts slightly lower production rates for acetate in conditions with higher amounts of acetate in the background.

**Figure 2:**
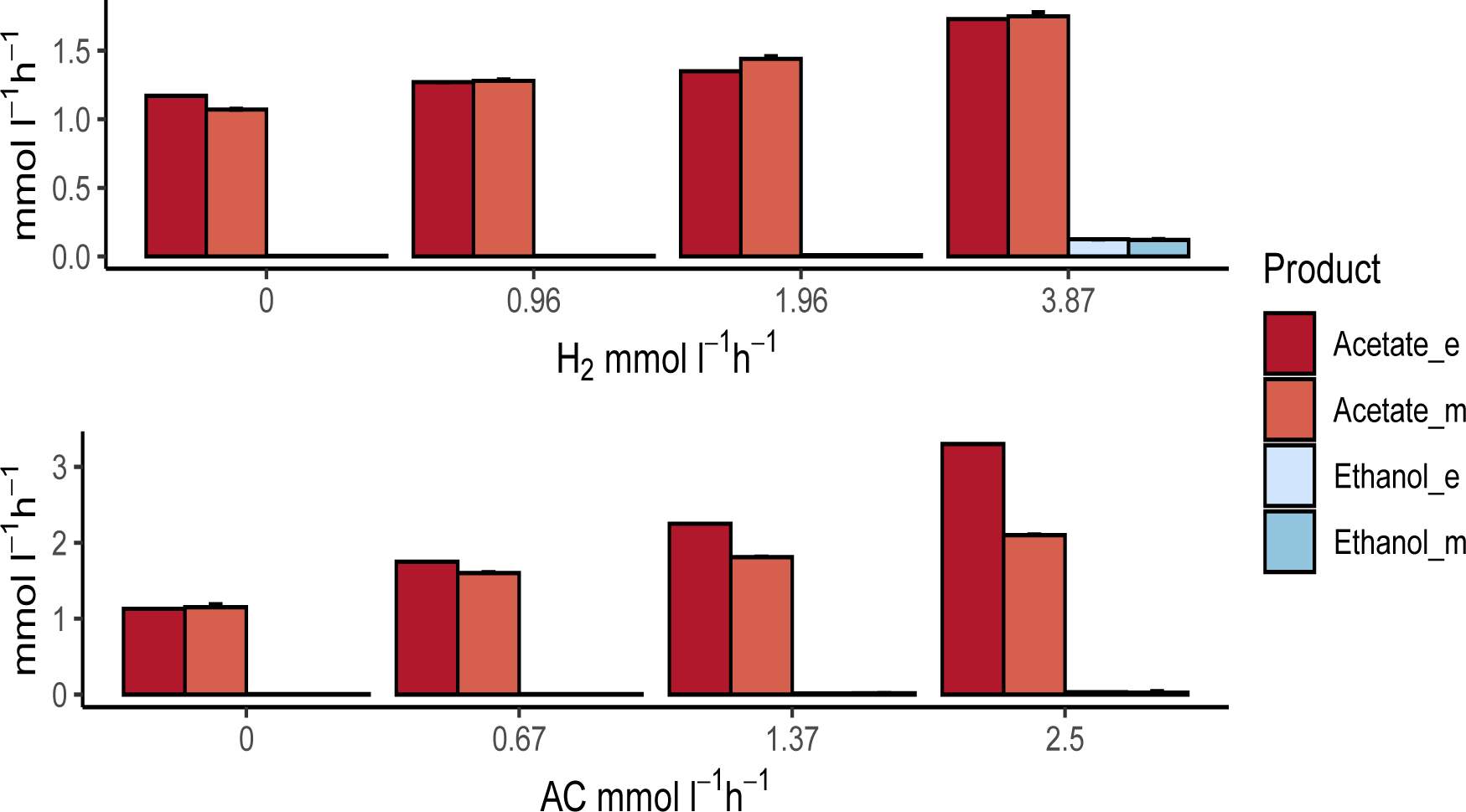
Comparison of the experimental (_e) and model (_m) results of the volumetric production rates of fermentation products of the *C. autoethanogenum* mono-culture under CO/H_2_ and CO/AC (acetate) conditions. In CO/H_2_ conditions, CO feed rate= 4.8 mmol l^−1^ h^−1^; growth rate=0.021 h^−1^ and working volume= 1 l. In CO/AC conditions, CO feed rate= 6.4 mmol l^−1^ h^−1^; growth rate=0.028 h^−1^ and working volume= 0.75 l. Substrates feed rates and production rates are expressed in mmol l^−1^ h^−1^.

The co-culture experiments were run under same conditions as the mono-culture experiments. Figure 3 shows the comparison between experimental results collected in co-culture experiments [11] and the results obtained via the multi-species model. The model correctly predicts production of medium-chain fatty acids upon introduction of *C. kluyveri*. Similarly to the mono-culture simulations, there is a slight mismatch between predicted and observed acetate production as the model predict slightly higher acetate production rates than those measured. The model correctly predicts the increase of medium-chain fatty-acids when more H_2_ or acetate is added. Hexanoate increases gradually the more acetate or H_2_ is added in the media. When H_2_ feed rate is equal to 5.3 mmol l^−1^ h^−1^, butanol is also produced (0.075 mmol l^−1^ h^−1^). Also, ethanol accumulation is low in most co-culture conditions, suggesting most of it is metabolized by *C. kluyveri*.

**Figure 3:**
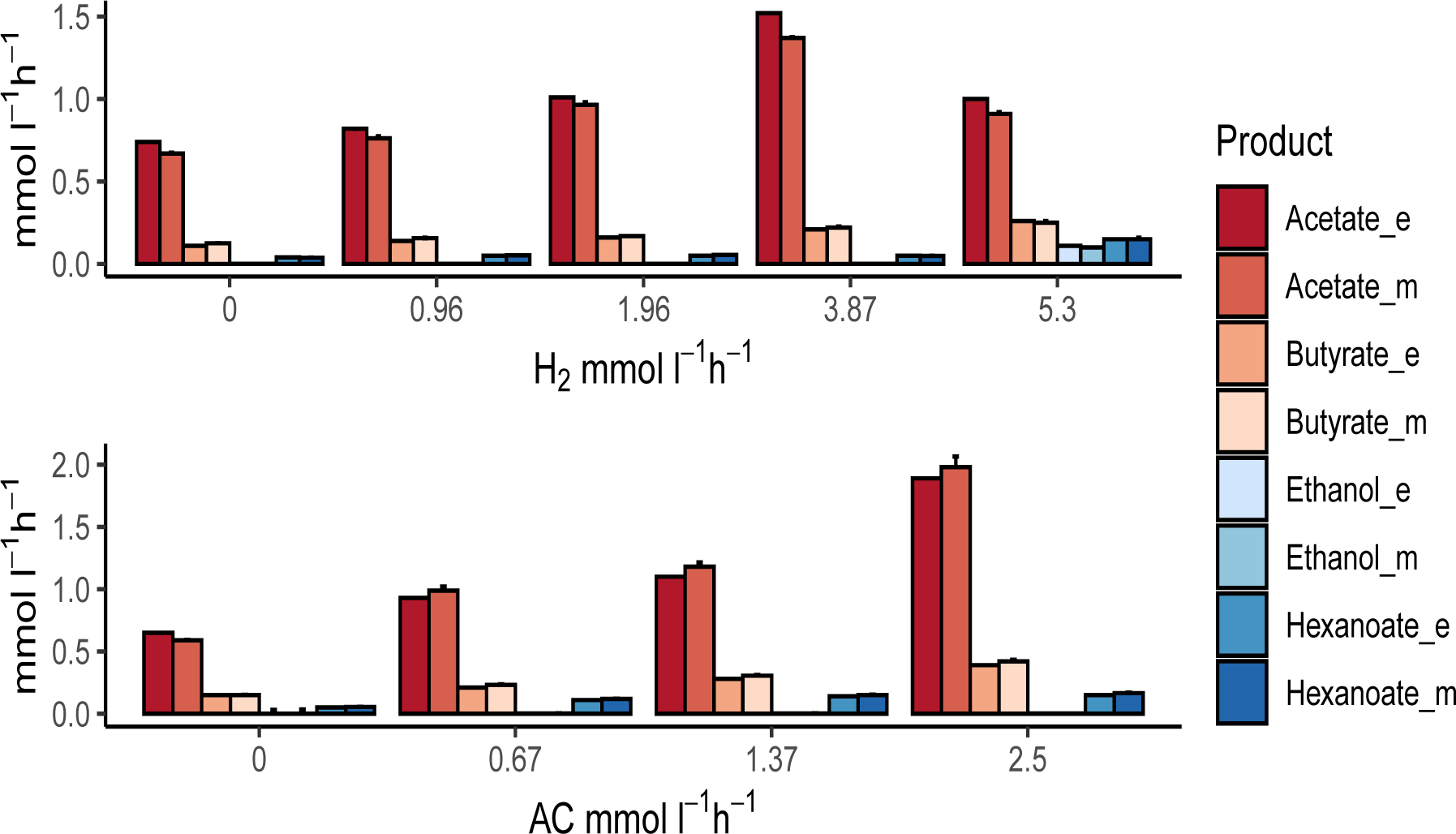
Comparison of the experimental (_e) and model (_m) results of the volumetric production rates of the fermentation products in co-culture under CO/H_2_ and CO/AC (acetate) conditions. In CO/H_2_ experiments, CO feed rate= 4.8 mmol l^−1^ h^−1^; growth rate=0.021 h^−1^ and working volume= 1 l. In CO/AC conditions, CO feed rate= 6.4 mmol l^−1^ h^−1^; growth rate=0.028 h^−1^ and working volume= 0.75 l. Feed rates of substrates and production rates are expressed in mmol l^−1^ h^−1^.

#### 3.3.1. Assessing the distribution of metabolic fluxes with the multi-species model

After having shown that the model describes accurately the metabolic interactions between the two microbes, we used it to explore intracellular flux distributions that would be otherwise challenging to access. To study the metabolic fluxes in the co-culture, we used a sampling approach that produces, for each reaction in the combined model, a distribution of possible fluxes. Figure 4 provides an overview of selected reactions in the system (indicated by R#). Fluxes for all reactions can be found in the supplementary material.

**Figure 4:**
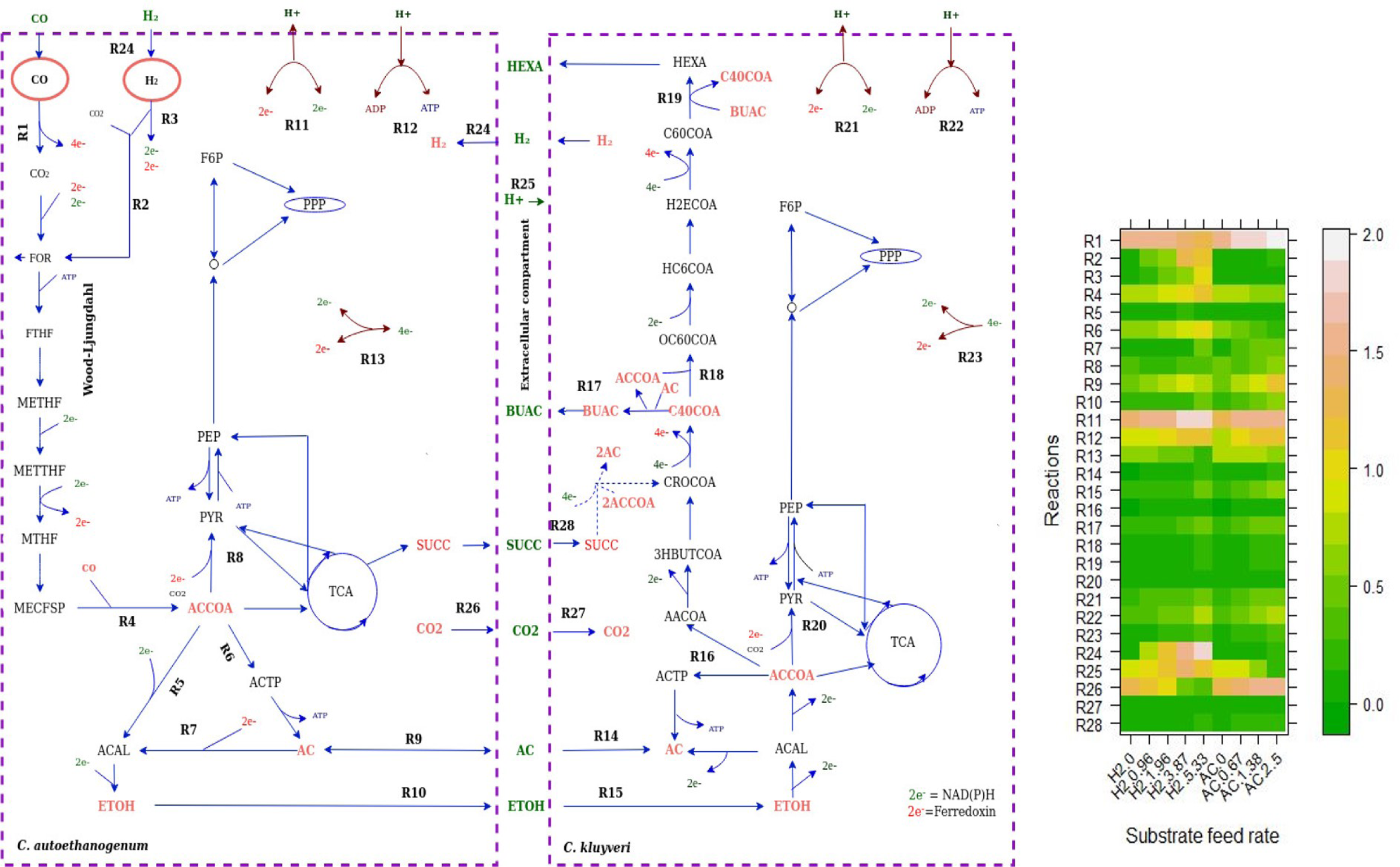
Schematic representation of the simulated metabolism of *C. autoethanogenum* and *C. kluyveri* under chemostat cultivation conditions. In CO/H_2_ experiments, CO feed rate= 4.8 mmol l^−1^ h^−1^; growth rate=0.021 h^−1^ and working volume= 1 l. In CO/AC conditions, CO feed rate= 6.4 mmol l^−1^ h^−1^; growth rate=0.028 h^−1^ and working volume= 0.75 l. Blue arrows indicate the fluxes direction. Metabolites colored green are extracellular metabolites, partly exchanged between species and partly excreted or assimilated to/from the media. The heatmap on the right shows the fluxes of the R# reactions selected on the map for all conditions. Flux values are log transformed (log(Flux+1)) for a better visualization

CO or CO and H_2_ are converted via the Wood-Ljungdahl pathway in *C. autoethanogenum*. In this pathway, CO is converted to CO_2_ via CO dehydrogenase, providing reducing equivalents to the cell. Released CO_2_ is shuttled into the Wood-Ljungdahl pathway via the bifurcating formate dehydrogenase [41]. H_2_ taken up by *C. autoethanogenum* is used for redox generation in NADP-dependent electron bifurcating hydrogenase (Hyt) reaction (R3) and in formate hydrogen lyase reaction (R2 in figure 4) to produce formate. The flux through these two reactions increases with increasing H_2_ supply. Part of the formate is excreted and part is further metabolized to acetyl-CoA (AC-COA) following the Wood-Ljungdahl pathway. Pyruvate is partly produced from acetyl-CoA via the pyruvate synthase (R8 in figure 4) for assimilation. The majority of the acetyl-CoA is converted to acetate via acetyltransferase (R6) and acetate kinase, yielding ATP. Ethanol can be formed in two ways [42]: from the reduction of acetate to ethanol via aldehyde ferredoxin oxidoreductase (R7) and alcohol dehydrogenase, or via reduction of acetyl-CoA to acetaldehyde (ACAL) and ethanol. Acetate and ethanol are secreted to the medium where it is partly taken up by *C. kluyveri*. In *C. kluyveri* ethanol is oxidized to acetyl-CoA (ACCOA) and part of acetyl-CoA is converted to acetate via acetyltransferase and acetate kinase (R16). Acetyl-CoA initiates the reverse *β*-oxidation pathway (R16-R19) to produce butyrate via acetoacetyl-CoA (AACCOA), then 3-hydroxybutyryl-coa (3HBUTCOA), crotonyl-CoA (CROCOA) and butyryl-CoA (C40COA). C40COA, transfers the CoA group to acetate, producing butyrate and acetyl-CoA. Some of the butyrate can be elongated further to hexanoate by reaction of butyryl-CoA together with hexanoyl-CoA (C60COA)(R19). acetyl-CoA is also assimilated by fixing CO_2_ to pyruvate via pyruvate synthase (R20) in *C. kluyveri*.

The model indicates that part of the acetate pool in *C. kluyveri* comes from uptake of succinate produced by *C. autoethanogenum* (see R28). Succinate is converted to Crotonyl-CoA (CROCOA), yielding an additional 2 acetate (see figure 4, involving the pathway via succinyl-CoA, succinate semi-aldehyde, 4-hydroxybutyrate, and 4-hydroxybutyryl-CoA [32]. The model predicts a low amount of ethanol being oxidized to acetate (R16), supporting activity of the succinate pathway.

The model predicts reduction of CO_2_ production by *C. autoethanogenum* with increasing H_2_ feed rate (figure 4B). This was also observed in the experimental measurements [11], where CO_2_/CO ratio decreased linearly with increasing hydrogen uptake. According to model predictions, CO_2_ production rate drops to 0.28 mmol l^−1^ h^−1^ when H_2_ is supplied, as compared to 2.5 mmol l^−1^ h^−1^ that is produced when CO is the only carbon source (see supplementary material). It is observed that more H_2_ is metabolized by *C. autoethanogenum* with increasing H_2_ feed rate (as shown in figure 4B) similar to what was found in experimental results [11]. When more H_2_ is fed to the reactor (see R25), more protons are released. In contrast, less protons are released when more acetate is fed. ATP synthase increases in both species (R12, R22) when more H_2_ or acetate is fed to the reactor.

### 3.4. Effect of the biomass ratio of C. kluyveri-C. autoethanogenum on the metabolic profiles of the culture

The model enables detailed inspection of production and uptake profiles. Therefore we investigated the sharing of metabolites between both organisms. Analysis of intracellular fluxes in figure 4 suggests that succinate is produced by *C. autoethanogenum* and metabolized by *C. kluyveri* producing part of the acetate and crotonyl-CoA pool needed for chain elongation and production of fatty-acids.

Figure 5 shows the different profiles depending on the biomass species ratio when we simulate chemostat cultivation experiments of *C. autoethanogenum* and *C. kluyveri* growing on CO and H_2_ as carbon and energy sources. Succinate, acetate and ethanol uptake by *C. kluyveri* increases with more H_2_ supply. Succinate uptake decreases when *C. kluyveri* is less abundant in the co-culture. On the contrary, the amount of ethanol and acetate metabolized by *C. kluyveri* decreases if *C. kluyveri* is more abundant in the co-culture.

**Figure 5:**
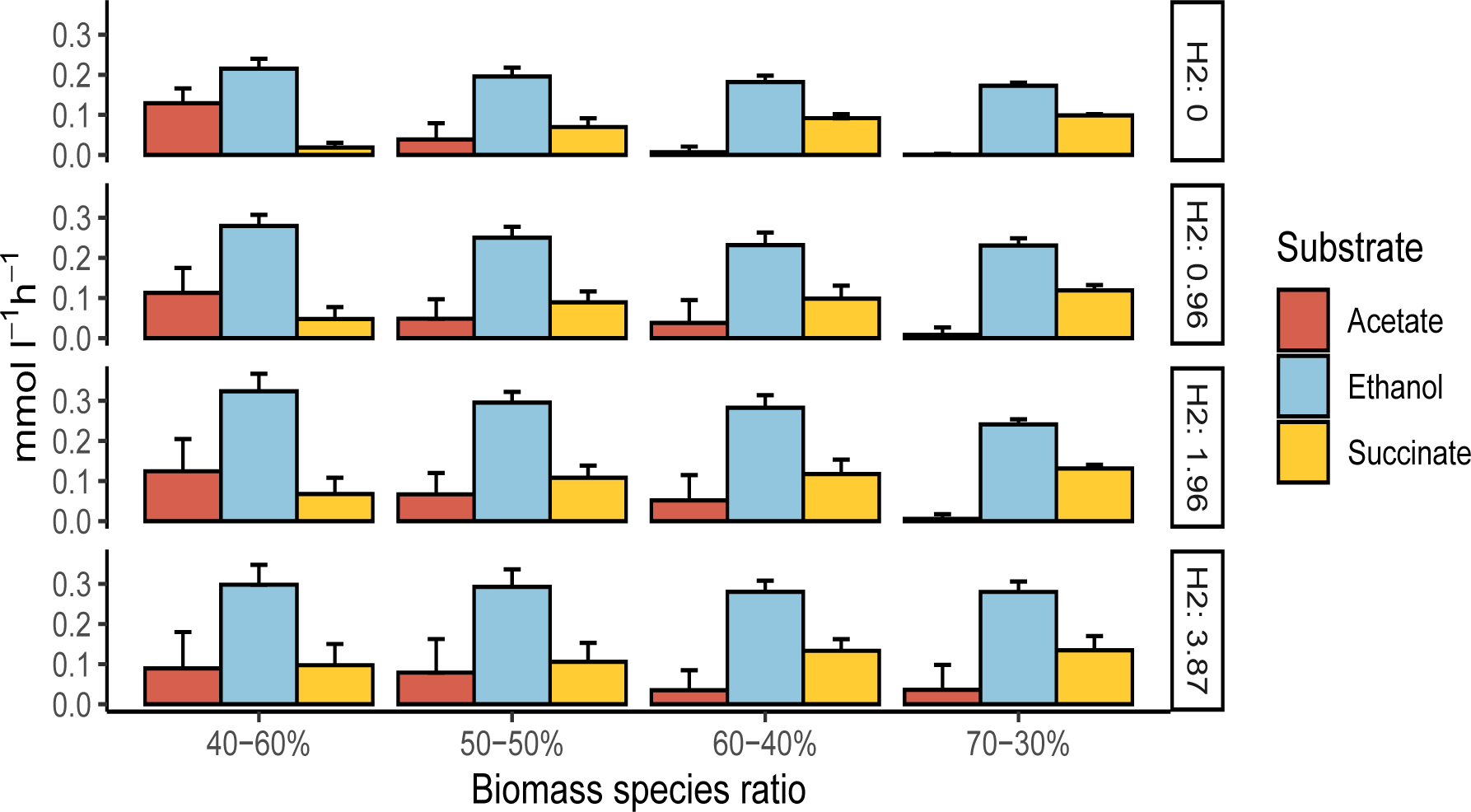
The effect of changing biomass species ratios on the uptake of acetate, ethanol and succinate by *C. kluyveri* at different H_2_ feed rates and fixed CO feed rate. CO= 4.8 mmol l^−1^ h^−1^ and growth rate=0.021 h^−1^, respectively. *C. kluyveri* -*C. autoethanogenum* biomass ratio are indicated on the x axis and y axis represents the fluxes of transport reactions from extracellular to *C. kluyveri* compartment of acetate, ethanol and succinate.

To further investigate the role of succinate, similar simulations were performed but this time preventing the uptake of succinate by *C. kluyveri* (figure 6). The model shows that, without succinate uptake, a biomass-species ratio of 70-30% and CO as the only carbon source results in an unfeasible situation. In addition, the fluxes through reactions related to non-growth associated maintenance (ATPM, ATPM auto) decreased substantially when biomass ratios of 60-40%, 50-50% and 40-60% were considered. Thus, the *in silico* analyses shows that the only way to meet the experimentally observed constraints is through succinate uptake. Changes in the biomass ratio also affect acetate and ethanol exchange between the microbes. Acetate and ethanol uptake by *C. kluyveri* increase when more H_2_ is fed to the system. Acetate/ethanol production ratios become higher when *C. kluyveri* is more abundant and decrease when there is more *C autoethanogenum* in the co-culture. Based on these results, the species biomass ratio in co-cultivation is estimated to be between 60-40% and 70−30% (*C. kluyveri*−*C. autoethanogenum*).

**Figure 6:**
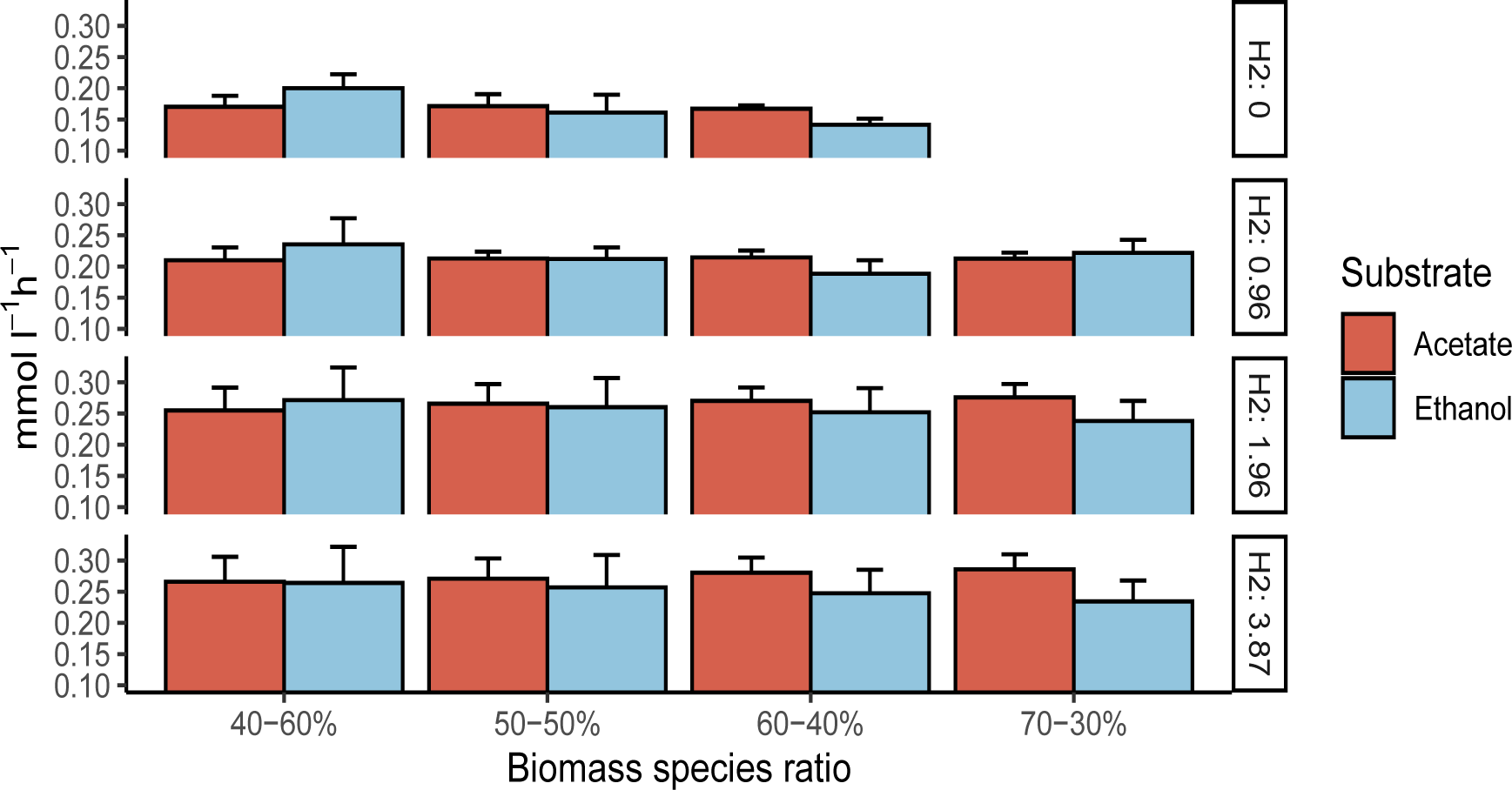
Effect of changing biomass species ratios on the uptake of acetate and ethanol by *C. kluyveri* at different H_2_ feed rates without succinate uptake and fixed CO feed rate. CO= 4.8 mmol l^−1^ h^−1^ and growth rate=0.021 h^−1^.*C. kluyveri* − *C. autoethanogenum* biomass ratio are indicated on the x axis and y axis represents the fluxes of transport reactions from extracellular to *C. kluyveri* compartment of acetate and ethanol.

### 3.5. Strategies to increase production of medium-chain fatty-acids

We used the experimentally-validated model to simulate alternative scenarios that have so far not been explored experimentally. Accordingly, we present here an analysis of possible strategies to increase medium-chain fatty acid production by the co-culture.

#### 3.5.1. Addition of succinate

As a strategy to increase the production of desired products, we simulated how the addition of succinate as extra carbon source would affect the production of butyrate and hexanoate at different H_2_ feed rates and fixed CO feed rate (4.8 mmol l^−1^ h^−1^). Figure 7 shows the product profile for the species biomass ratio 70 − 30% (*C. kluyveri* -*C. autoethanogenum*), respectively. The increased carbon availability has been considered and used to normalize the results, so they are represented as mmol of product per total substrate (CO and succinate) per carbon.

**Figure 7:**
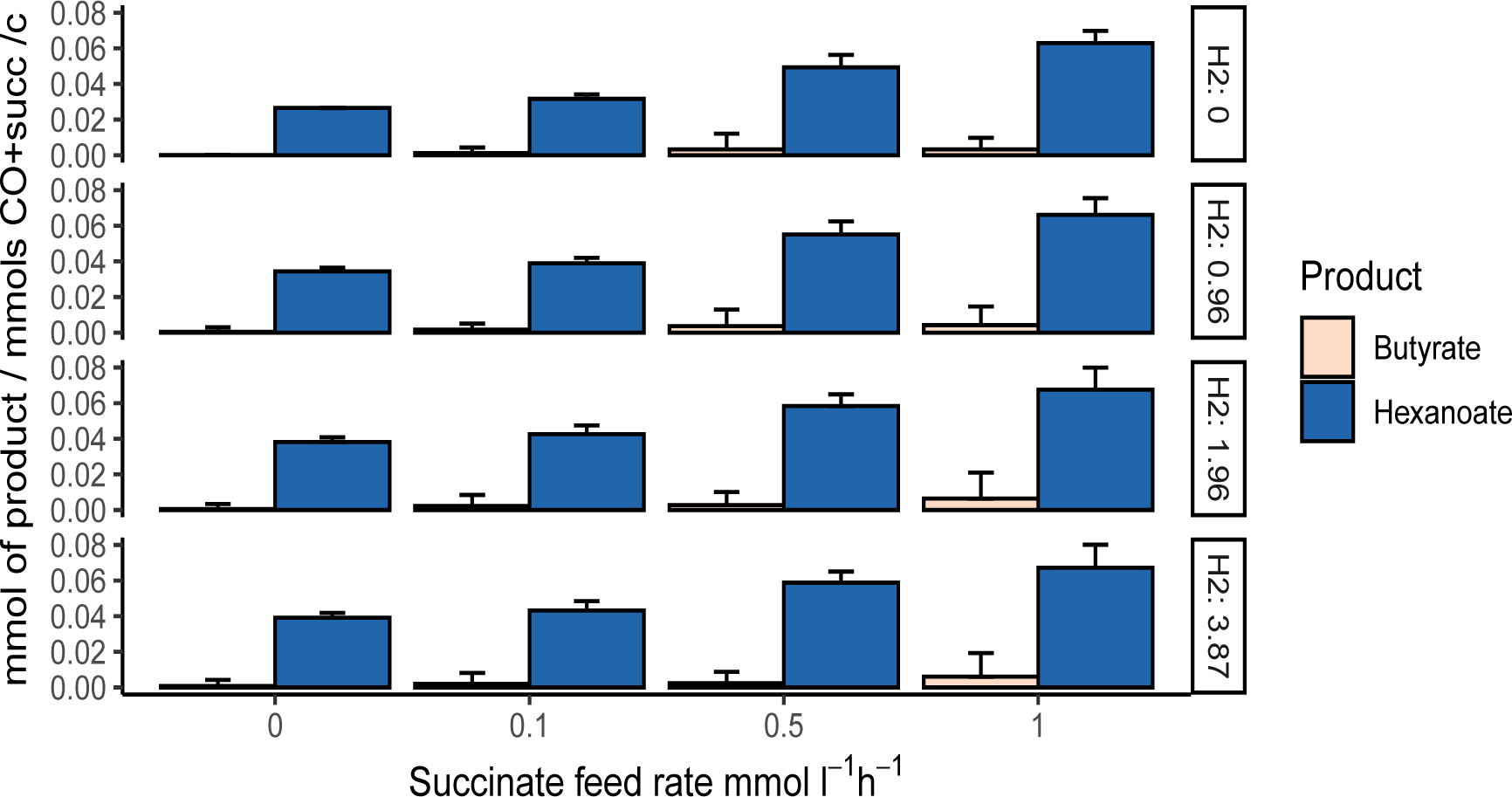
Effect of succinate addition on the production of butyrate and hexanoate under different H_2_ feed rates when a biomass ratio of 70 − 30% is considered. X-axis represent succinate feed rate and y-axis represent mmol of butyrate or hexanoate normalized per mmols of total substrate per carbon.

As already indicated in figure 3, and in contrast to experimental results, the model predicts more hexanoate than butyrate production even when succinate uptake is 0. The model predicts a yield of 0.026 mmol of hexanoate per mmol of CO. This means, that this co-culture has the capacity to produce three times more hexanoate than what it is currently being produced (0.009 mmol per mmol of CO) [11], reducing the gap respect to the maximum theoretical yield (0.056 mmol of hexanoate per mmol of CO) to 3%. Increased hexanoate production is observed next to an increase in CO_2_ uptake by *C. kluyveri* and changes in H+ balance and ATP synthase.

Furthermore, succinate addition allows increased production of both fatty-acids. Hexanoate production can increase up to four times when succinate is added and butyrate has potential to increase around five times with respect to the results obtained with no succinate addition. On the other hand, simulations show no relevant differences upon variations of H_2_ feed rates.

#### 3.5.2. Genetic intervention strategies

As a second strategy for co-culture optimization we have predicted and evaluated genetic interventions that could lead to higher medium-chain fatty-acid production.

The strain design algorithms OptKnock and RobustKnock [39, 40] were applied to identify candidate reactions to be knocked out. These were subsequently evaluated through dedicated simulations. Figure 8 shows the effect of knocking out three candidate reactions on the production of ethanol in *C. autoethanogenum*. The three reactions are located in the metabolism of *C. autoethanogenum*. Mutant 1 refers to the knock out of formate transport reaction via proton symport (FORt2, model id rxn05559_c0). Mutant 2 refers to the deletion of acetaldehyde:NAD+ oxidoreductase (ACALDx, model id rxn00171_c0). Mutant 3 refers to the knock out of formate dehydrogenase (ferredoxin) (FDH fer, model id rxn00103_c0). The impact of removal of these reactions is compared to wild type *C. autoethanogenum*, corresponding to the GEM without any modification. The deletion of each reaction results in increased ethanol production. Ethanol increases up to 83% with respect to the wild type in simulations with CO as the only carbon source. In conditions where H_2_ acts as second substrate, ethanol production increases up to 150%. Acetate decreases up to 11% in simulations with only CO and decreases up to 30% in CO/H_2_ simulations. Mutant 1 seems to have a higher ethanol yield when CO is the only carbon source compare to CO/H_2_ conditions while mutant 2 and mutant 3 provide a higher effect on ethanol/acetate ratio in CO/H_2_ simulations. As seen in Figure 6, the production of ethanol by *C. autoethanogenum* and therefore, the uptake of ethanol by *C. kluyveri*, increases with increasing H_2_ feed rate, resulting in an increased production of medium-chain fatty-acids. This suggests that these deletions in *C. autoethanogenum* can improve the production of medium-chain fatty-acids in this respective co-culture, since ethanol production can potentially be increased.

**Figure 8:**
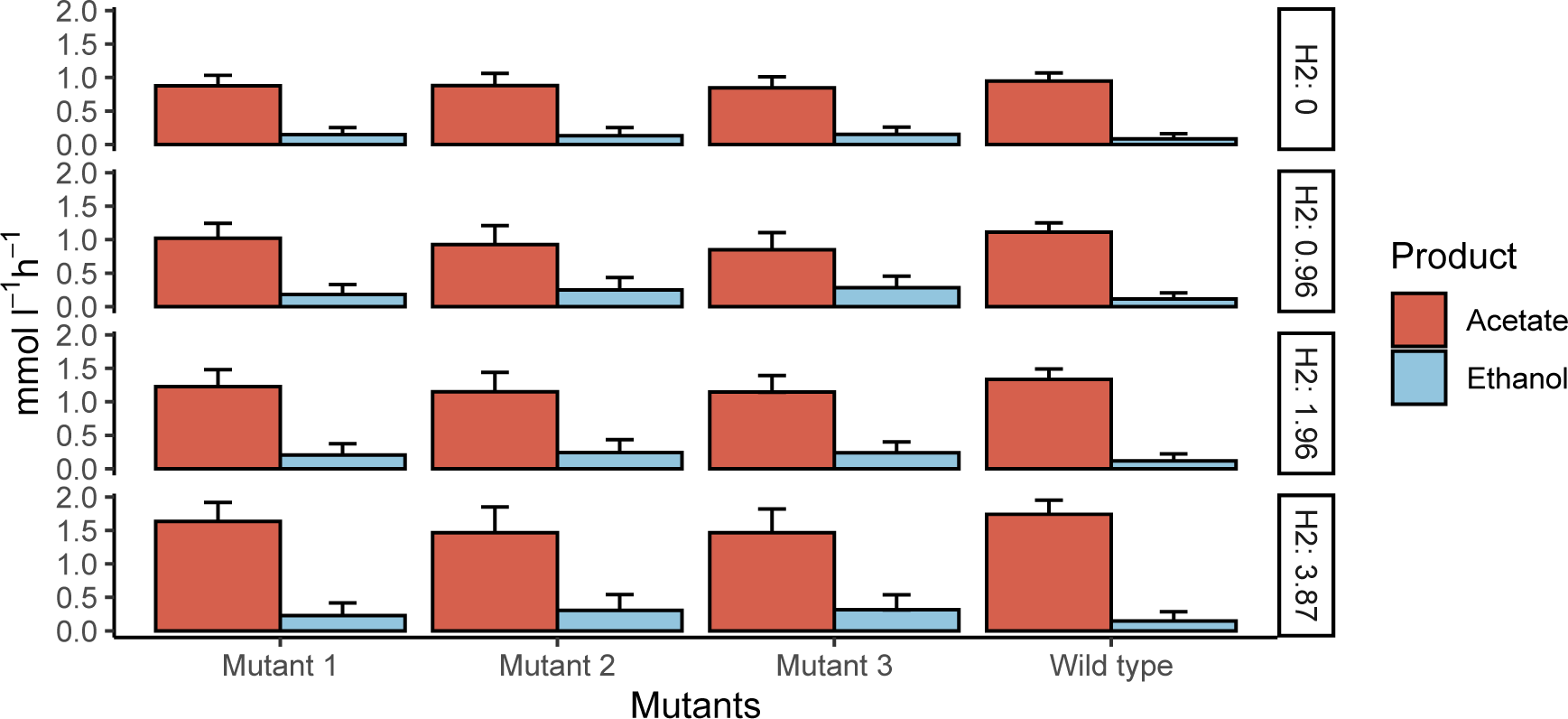
Effect of single reactions deletion on the production of ethanol in *C. autoethanogenum*. X axis represents the mutants applied on the GEM of *C. autoethanogenum* and y axis represents the production of acetate and ethanol. Mutant 1: Formate transport in via proton symport (rxn05559_c0). Mutant 2: acetaldehyde:NAD oxidoreductase (rxn00171_c0); Mutant 3: formate dehydrogenase (ferredoxin) (rxn00103_c0)

## 4. Discussion

We present here a constraint-based model of the co-culture of *C. autoethanogenum* and *C. kluyveri* in the context of CO/syngas fermentation to produce medium-chain fatty-acids. A model with similar characteristics had already been used to simulate CO to butyrate conversion by bacterial co-culture systems [34]. Our model extends previous efforts, and was calibrated and tested with a battery of available experimental measurements. Modeling bacterial communities using flux balance analysis and GEM is complicated by the fact that special attention has to be paid to the biomass abundances of the microbial species in order to achieve balanced growth of the co-culture. Previous efforts used dynamic flux balance analysis to consider biomass growth [34]. Here, steady-state conditions were used of which additional data was available, allowing to overcome the challenge of estimating relative abundance of each species by combining microscopy observations and RNA seq in terms of cell numbers. These were subsequently converted to biomass ratios by considering the relationships between cellular dimensions, cell volume and biomass dry weight. This enabled the application of community flux balance analysis [26], which has been shown to accurately predict flux distributions and exchange fluxes between species and the community environment when analyzing stable communities in chemostat experiments.

The presented co-culture model is versatile and can simulate the CO/syngas fermentation process leading to medium-chain fatty-acid production. The presented model describes both the behaviour of a *C.autoethanogenum* mono-culture thriving on syngas as well as the behaviour of a co-culture of *C. autoethanogenum* and *C.kluyveri*. The model accurately reproduces the volumentric production rates of fermentation products obtained in chemostat experiments, and predicts the shift of the metabolism of *C. autoethanogenum* towards solventogenesis [11]. In addition, model predictions on production/consumption rates (see figure 4) agree reasonably with previous literature on CO/syngas fermentation [32, 12, 10, 11].

Analyses of metabolic fluxes in the model suggested succinate production by *C. autoethanogenum* as an important intermediate in the co-culture. Accumulation of succinate has not been experimentally observed in the calibration experiments [11], and is not described as major physiological end-product of *C. autoethanogenum* when grown on syngas [12]. However, its production by *C. autoethanogenum* in the presence of *C. kluyveri* seems possible as it has been reported to be an overflow product of acetogenic metabolism [43]. Here, succinate is produced to overcome the temporal overflow of C/electrons, potentially in conditions where too much reduction equivalents are provided. Succinate is described as a possible substrate for *C. kluyveri* [32]. Presence of succinate could slow down consumption of ethanol/acetate by *C. kluyveri*. This would also affect co-culture compositions by limiting *C. kluyveri* abundances.

The model predicts succinate to be an important intermediate in the co-culture, as omitting its uptake resulted in unfeasible growth conditions when grown on only CO (see figure 6, ratio 70-30%). This dependency is additionally shown in the case with both CO and H_2_, where ATPM decreased substantially to sustain growth. An alternative explanation for the succinate dependency could be related to *C. autoethanogenum* cell size assumptions, being potentially bigger than the average size, since it was reported to have considerable variations [12]. Therefore, the co-culture might operate in ratios where the biomass of *C. autoethanogenum* is more abundant, with relative values between 60-40% and 70 − 30%.

As the model closely predicted obtained experimental values, it can be used to design potential strategies for improved production. Here, we explored a series of strategies to optimize the production of hexanoate. The first strategy relied on the role of succinate, as the model predicts it increases the pool of acetate and crotonyl-CoA, a precursor of the desired fatty-acids, in *C. kluyveri* (see figure 4). The flux through the reverse *β*-oxidation pathway increases up to four times when succinate is added (see supplementary material). The increase of crotonyl-CoA results in a higher butyryl-CoA pool. The presence of more butyryl-CoA initiates the chain-elongation process to produce hexanoyl-CoA in the same way as butyryl-CoA is formed (see 4). According to the model, most of butyrate formed from butyryl-CoA reacts with hexanoyl-CoA producing hexanoate. This results in an increase in hexanoate production up to three times (see figure 7). Moreover, an increase in ethanol uptake by *C. kluyveri* and a decrease in acetate production by *C. autoethanogenum* and subsequent uptake by *C. kluyveri* appears to cause an additional boost in hexanoate production. According to the model, addiction of succinate addition raise hexanoate production up to 0.067 mmol per carbon of fed substrate (CO and succinate) and would possibly lead to a further increased production of MCFA and alcohols in conditions with higher H_2_ influx (*>* 5 mmol l^−1^ h^−1^), as it has been observed previously.

The second strategy aims to increase hexanoate by increasing ethanol production by *C. autoethanogenum*. An increasing ratio of ethanol/acetate ratio has been shown to result in increased hexanoate production in *C. kluyveri* [44, 35]. In cases where butyrate and hexanoate are not constrained (see figure 7), the predicted ethanol/acetate ratio (around 6:4) is higher than when butyrate is more prominent (see figure 6). The model thus confirms previous experimental results and highlights the potential of an increased ethanol/acetate ratio in stimulating the production of hexanoate. It should be bear in mind that model predictions are based on optimality principles and assumptions. The model suggests that increased ethanol production leads to increased medium-chain fatty-acids. Still, data regarding the impact of *C. kluyvery* on the behaviour of the co-culture, would be needed for accurate predictions.

*C. autoethanogenum* is known to increase production of ethanol under acidic or redox overloading conditions [45, 46, 11]. Additionally, partial inactivation of the adhE cluster or knockout of one of the AOR genes has been shown to result in increased ethanol production in *C. autoethanogenum* [42]. Using the GEM herein developed for *C. autoethanogenum* we predicted that the individual knock-out of one of the following three reactions could increase the production of ethanol: acetaldehyde oxidoreductase (ACALDx), formate transport (FORt2), and the bifurcating formate dehydrogenase (ferredoxin) (FDH fer) (see figure 8). The acetaldehyde oxidoreductase reaction (ACALDx, id rxn00171_c0) is associated with several isoenzymes encoded by genes: CAETHG-RS16140, CAETHG-RS08865, CAETHG-RS08810, CAETHG-RS18400 and CAETHG-RS18395. The same isoforms are associated to aldehyde ferredoxin oxidoreductase reaction (CODH-ACS) (leq000004) and two of them (CAETHG-RS18400 and CAETHG-RS18395) are also involved in ethanol oxidoreductase (ALCDx) (rxn00543_c0) reaction. The affinity of each isoenzyme to each reaction has to be studied in order to fully eliminate acetaldehyde oxidoreductase activity. An acetaldehyde oxidoreductase (ACALDx) mutant has previously been shown to indeed have increased ethanol production up to 180% [42], making the deletion of this reaction seems a promising application to increase fatty-acids production in the co-culture system. The knock out of formate-related reactions in *C. autoethanogenum* is not described previously, but model simulations done here, suggest that they contribute to ethanol production. Formate transport in via proton symport (FORt2, model id rxn05559_c0) is catalyzed by an enzyme encoded by only one gene -CAETHG-1601, which allows relatively easy removal of this activity. The model predicts that inactivation of this reaction forces more flux through the Wood-Ljungdahl pathway, increasing the amount of acetyl-CoA. Due to the increase in acetyl-CoA pool, the fluxes through aldehyde ferredoxin oxidoreductase and acetaldehyde oxidoreductase are increased, thus producing more ethanol. As third option the model suggests to knock out the formate dehydrogenase (ferredoxin) (FDH fer, model id rxn00103_c0) activity. This reaction has three associated isoenzymes encoded by genes: CAETHG-RS00400, CAETHG-RS13720 and CAETHG-RS14690. The inactivation of this reaction forces the production of formate mostly via formate hydrogen-lyase from H_2_ (FHL, rxn08518_c0). In mono-culture conditions where there is no H_2_ supply, H_2_ is produced via an NADP-dependent electron-bifurcating hydrogenase reaction (Hyt) (leq000001). This functionality of Hyt seems to occur in situations where redox mediators get too reduced [41]. The model shows a higher conversion of CO converted to CO_2_ via CO dehydrogenase (CODH4, model id rxn07189_c0) which forces more flux through Wood-Ljungdahl pathway, producing more acetyl-CoA. Also, more CO_2_ is fixed producing pyruvate via pyruvate synthase (rxn05938_c0) which is shuttled back via pyruvate formate lyase (rxn00157 Cc0) to produce more acetyl-CoA and formate. The extra pool of acetyl-CoA forces more flux through aldehyde ferredoxin oxidoreductase and acetaldehyde oxidoreductase which leads to more ethanol.

When the reactions of fatty-acids production are not constrained, the model always predicts more hexanoate as compared to butyrate production measured in the actual chemostat experiments [11]. A potential reason for this is the pH of the co-culture and related toxicity effects of medium-chain fatty acids. Hexanoate production in *C. kluyveri* has been reported to be better at higher pH [47]. Thus, potentially the pH of 6.2 in the co-culture limits its production in the actual experiments. In addition to toxicity effects, the function of membrane proteins such as ATP synthase, electron transport chains or transporters can be affected by the change of proton motive force at different pH [48]. This is reflected by the observation that model predictions show differences in ATP synthase and proton balance under different butyrate/hexanoate production conditions (see supplementary material and figure 4). However, the acid stress response is difficult to simulate in GEMs, which possibly results in the differences observed between the model prediction and experimental results.

We observe an increase of CO_2_ uptake by *C. kluyveri* in cases where hexanoate is more abundant (see figure 4 and supplementary material) compared to simulations of experimental conditions, where butyrate is more abundant (see figure 4). CO_2_ uptake is essential for growth and C1 intermediate production in *C. kluyveri* [49]. Model predictions show that around 60% of CO_2_ is metabolized following phosphoenolpyruvate carboxylase reaction (PPC), where CO_2_ is fixed together with phosphoenolpyruvate (pep) producing oxaloacetate (oaa). Oaa produces aspartate via aspartate aminotransferase (Rckl310). Aspartate is a precursor in the synthesis of threonine involving aspartate kinase (Rckl334), aspartate semialdehyde oxidoreductase (Rckl323), homoserine kinase (Rckl335) and threonine synthase (Rckl336). Then, threonine produces acetaldehyde and glycine via threonine aldolase (Rckl341). Acetaldehyde is converted to acetyl-CoA following aldehyde alcohol dehydrogenase (ADH) reaction. So, the increase in CO_2_ assimilation could lead to more acetyl-CoA and thus, more fatty-acids. Around 30-40% of CO_2_ is converted to carbonic acid. Carbonic acid produces oxaloacetate via pyruvate carboxylase (Rckl014), which follows the aforementioned route to acetyl-CoA. The CO_2_ left is fixed with acetyl-CoA in pyruvate synthase (Rckl119) to produce pyruvate that produces again acetyl-CoA and formate via pyruvate formate lyase (PFL). Formate is assimilated following the tetrahydrofolate pathway producing C1 compounds such as serine, glycine and tetrahydrofolate. Simulations made with higher CO_2_ uptake rates than the ones predicted when hexanoate is more abundant (*>*0.75 mmol l^−1^ h^−1^) however, did not lead to higher production rates of hexanoate or butyrate. This suggest that *C. kluyveri* metabolizes CO_2_ up to a maximum value. In fact, this is supported by the observed correlation between growth and the maximum CO_2_ fixed by *C. kluyveri* [50]. The use of lower hydraulic retention time and high pressure bioreactors, could possibly increase the uptake of CO_2_, close to its maximum capacity.

The maximum hexanoate predicted by the multi-species GEM is reached when succinate is added into the system in combination with CO and H_2_ (see figure 7). Table 1, shows a comparison of electron yields for the hexanoate production predicted in this study compared to hexanoate production in other studies with similar culture systems [51, 52, 11]. Electron yield is calculated based on the amount of electrons going to hexanoate per the total amount of electrons going into the system as carbon and energy sources. Co-cultures of *C. autoethanogenum* and *C. kluyveri* yielded more hexanoate growing on CO/H_2_ or CO and acetate compared to a co-culture of *C. ljundahlii* and *C. kluyveri*, potentially as in the latter relatively more alcohols and C8 acids were produced as well. A mixed culture enriched in *Acinetobacter, Alcaligenes*, and Rhodobacteraceae growing solely on CO [52], increased the electron yield with respect to previously mentioned co-cultures up to 0.32. However, the addition of succinate in co-cultivation of *C. autoethanogenum* and *C. kluyveri* grown on CO and H_2_ (this study), is here predicted to increase the yield of hexanoate up to 0.6, reflecting the potential of this approach to produce medium-chain fatty-acids.

**Table 1:** Comparison of electron yield obtained for hexanoate production between predicted results by the multi-species GEM and other coculture/mixed culture. Electron yield is expressed by the amount of electrons going to hexanoate per total amount of electrons entering the system.

|  | Substrates <sup>a</sup> | Hexanoate <sup>b</sup> | Electron yield <sup>c</sup> |
| --- | --- | --- | --- |
| *C. ljungdahlii* & *C. kluyveri* [51] | CO=31.9<br>H<sub>2</sub>= 79.1 | 0.48 | 0.07 |
| Mixed culture [52] | CO=0.58 | 0.011 | 0.32 |
| *C. autoethanogenum* & *C. kluyveri* [11] | CO=4.8 <sup>d</sup><br>H<sub>2</sub>=5.3 | 0.15 | 0.26 |
| *C. autoethanogenum* & *C. kluyveri* [11] | CO=6.5 <sup>d</sup><br>AC=2.5 | 0.15 | 0.15 |
| Multi-species GEM | CO=4.8<br>H<sub>2</sub>=3.87<br>SUCC=1 | 0.59 | 0.6 |

<sup>a</sup>Volumetric consumption rate mmol l<sup>-1</sup> h<sup>-1</sup>
<sup>b</sup>Volumetric production rate mmol l<sup>-1</sup> h<sup>-1</sup>
<sup>c</sup>hexanoate e<sup>-</sup>/total e<sup>-</sup> in ; CO= 2 e<sup>-</sup> per mol, H<sub>2</sub> = 2 e<sup>-</sup> per mol; AC(acetate)= 8 e<sup>-</sup> per mol; SUCC (succinate)= 14 e<sup>-</sup> per mol and hexanoate = 32 e<sup>-</sup> per mol
<sup>d</sup> Assuming 90% gas consumption

Table 1 shows a comparison of the maximum production of hexanoate found on several studies with similar cultures.

## 5. Conclusions

The generation of the multi-species GEM of *C. autoethanogenum* and *C. kluyveri* has provided insights into the fermentation of CO/syngas to medium-chain fatty acids by this co-culture. The prediction of intracellular flux distribution in this consortium enabled to uncover the potential importance of succinate uptake via *C. kluyveri* to produce butyrate, and suggested an effect of the biomass species ratio on the substrate profile of *C. kluyveri*. Simulations indicated that succinate addition might result in a substantial increase in hexanoate yield from syngas. In addition, the model of *C. autoethanogenum* shows that the deletion of reactions FORt2 or ACALDx or FDH fer in *C. autoethanogenum* potentially increase ethanol production, suggesting a potential increase in hexanoate production when these deletions were to be applied in co-culture experiments. Altogether, our model-driven approach has set a good basis for the systematic design of strategies to modulate and optimize the production of valuable chemicals from syngas.

## Acknowledgements

The research leading to these results has received funding from the Netherlands Science Foundation (NWO) under the Programme ‘Closed Cycles’ (Project nr. ALWGK.2016.029) and the Netherlands Ministry of Education, Culture and Science under the Gravitation Grant nr. 024.002.002.

## Author contributions

Sara Benito-Vaquerizo: Conceptualization, Methodology, Software, Formal analysis, Investigation, Writing-Original draft preparation. Martijn Diender: Formal analysis, Investigation, Writing - Review & Editing. Ivette Parera Olm: Formal analysis, Investigation, Writing - Review & Editing. Peter J. Schaap: Conceptualization, Writing - Review & Editing. Vitor A.P. Martins dos Santos: Funding acquisition, Supervision, Writing-Reviewing & Editing. Diana Z. Sousa: Project administration, Funding acquisition, Conceptualization, Writing-Reviewing & Editing. Maria Suarez-Diez: Supervision, Conceptualization, Methodology, Visualization, Writing-Reviewing & Editing.

**Appendix A. Supplementary data**

